# Surfing beta burst waveforms to improve motor imagery-based BCI

**DOI:** 10.1101/2024.07.18.604064

**Authors:** S. Papadopoulos, L. Darmet, M.J. Szul, M. Congedo, J.J. Bonaiuto, J. Mattout

## Abstract

Our understanding of motor-related, macroscale brain processes has been significantly shaped by the description of the event-related desynchronization (ERD) and synchronization (ERS) phenomena in the mu and beta frequency bands prior to, during and following movement. The demonstration of reproducible, spatially-and band-limited signal power changes has, consequently, attracted the interest of non invasive brain-computer interface (BCI) research for a long time. BCIs often rely on motor imagery (MI) experimental paradigms that are expected to generate brain signal modulations analogous to movement-related ERD and ERS. However, a number of recent neuroscience studies has questioned the nature of these phenomena. Beta band activity has been shown to occur, on a single-trial level, in short, transient and heterogeneous events termed bursts rather than sustained oscillations. In a previous study, we established that an analysis of hand MI binary classification tasks based on beta bursts can be superior to beta power in terms of classification score. In this article we elaborate on this idea, proposing a signal processing algorithm that is comparable to-and compatible with state-of-the-art techniques. Our pipeline filters brain recordings by convolving them with kernels extracted from beta bursts and then applies spatial filtering before classification. This data-driven filtering allowed for a simple and efficient analysis of signals from multiple sensors thus being suitable for online applications. By adopting a time-resolved decoding approach we explored MI dynamics and showed the specificity of the new classification features. In accordance with previous results, beta bursts improved classification performance compared to beta band power, while often increasing information transfer rate compared to state-of-the-art approaches.

**Significance statement:** Patterns of waveform-specific burst rate comprise an alternative, neurophysiology-informed way of analyzing beta band activity during motor imagery (MI) tasks. By testing this method on multiple electroencephalography datasets and comparing its corresponding classification scores against those of conventional power-based features, this work demonstrates that brain-computer interface applications could benefit from utilizing beta burst activity. This activity gives access to a reliable decoding performance often requiring short recordings. As such, this study shows that waveform-specific beta burst rates encode information related to imagined (and presumably real) movements and serves as the first step for a real-time implementation of the proposed methodology.

## Introduction

Time-locked changes in induced power within specific frequency bands, originally described in a number of seminal studies in motor neuroscience (Pfurtscheller, 1981; Pfurtscheller and Berghold, 1989; Pfurtscheller and Lopes da Silva, 1999), have long influenced the way in which we interpret macroscale recordings of brain activity such as those provided by electroencephalography (EEG). These studies have revealed a gradual reduction in brain signal power during an ongoing movement or motor imagery (MI) task in the mu (∼8-12 Hz) (Neuper et al., 2006; Pfurtscheller et al., 2006, 1997; Pfurtscheller and Lopes da Silva, 1999) and beta (∼13-30 Hz) (Pfurtscheller et al., 1997; Pfurtscheller and Lopes da Silva, 1999) frequency bands relative to baseline activity. This phenomenon is termed event-related desynchronization (ERD). The same studies have, moreover, described a relative-to-baseline increase in power in the beta band shortly following the end of the movement or MI (Alayrangues et al., 2019; Neuper et al., 2006; Pfurtscheller et al., 1996), known as event-related synchronization (ERS). These phenomena are especially marked over cortical areas contralateral to the real or imagined movement (Kobler et al., 2020; Little et al., 2019; Makeig et al., 2000; Pfurtscheller and Berghold, 1989; Pfurtscheller and Neuper, 1997; Seeber et al., 2016; Zich et al., 2023) and their topographies approximately match the somatotopic organization of the sensorimotor cortices (Gordon et al., 2023; Natraj et al., 2022; Penfield and Rasmussen, 1950). Taken together, these observations have given rise to the hypothesis that the ERD is an indication of brain processes pertaining to movement preparation and execution while the ERS is an indication of processes related to movement completion (Kilavik et al., 2013).

Given the reproducibility of the spatial and frequency specificity of the ERD and ERS, these neural markers are often exploited by non-invasive BCI applications, especially those that are based on MI paradigms (Jayaram and Barachant, 2018; Tangermann et al., 2012). Such paradigms, designed to reproduce consistent time-locked signal modulations, normally rely on transforming the recordings in the time-frequency domain (TF) (Brodu et al., 2011; Bruns, 2004; Herman et al., 2008) and then applying spatial filtering, most commonly using the common spatial pattern algorithm (CSP) (Blankertz et al., 2008; Koles, 1991; Müller-Gerking et al., 1999). This chain of signal transformations is expected to increase signal-to-noise ratio by extracting signal power in specific time windows and frequency bands of interest, and also to maximize the spatial disparity among different MI classes (e.g. ”left” or “right” hand, or “feet”), thus improving classification results and/or allowing for decoding of multiple commands with distinct signal features (Lotte, 2014; Lotte et al., 2018).

Although the ERD and ERS are consistently observed across subjects and recording modalities, their nature is not clear. Based on the assumption of amplitude modulation of sustained oscillations, these patterns are the result of signal power averaging in the TF domain over multiple trials. However, converging evidence suggests that, on the contrary, beta band activity occurs in short events termed bursts (Coleman et al., 2024; Jones, 2016; Little et al., 2019; Lundqvist et al., 2024, 2016; Shin et al., 2017; Torrecillos et al., 2018; Wessel, 2020) on the single trial level, therefore questioning the functional role of ERD and ERS altogether, at least within the beta band. Beta burst rate has been shown to be more behaviorally relevant in motor processes (Enz et al., 2021; Hannah et al., 2020; Little et al., 2019; Rayson et al., 2023; Szul et al., 2023; Wessel, 2020) than averaged beta band power. Additionally, recent studies have shown that beta bursts are not a unitary phenomenon but rather constitute heterogeneous events (Szul et al., 2023) with different functions, alluded to by the differential modulation of their rate and shape depending on task conditions (Langford et al., 2023; Papadopoulos et al., 2024) or movement phase (Rayson et al., 2023; Szul et al., 2023). As such, beta bursts have the potential to be a more sensitive marker of brain processes during real or imagined movements on the single trial level.

To test this hypothesis, in a previous study we examined multiple open MI EEG datasets. We analyzed the activity of channels C3 and C4 during binary classification tasks of hand MI (“left” vs “right” hand) and demonstrated that the waveform-resolved beta burst rate is superior to beta band power changes in terms of classification (Papadopoulos et al., 2024). In this article we streamline our approach. We develop an algorithm that is computationally efficient and can analyze an arbitrary number of recorded signals thus being comparable to state-of-the-art techniques. Beta burst waveforms, whose rate is expected to be maximally modulated during the trial period compared to baseline, are identified in calibration data. These bursts are used as data-driven kernels that filter the signals from all recording channels in the time domain. The convolved signals are then spatially filtered with CSP and the spatial features are used as classification features. We re-analyze the activity during “left” and “right” hand MI of the same open EEG datasets and also a recently-published composite EEG dataset, now in a time-resolved fashion. We show that classification features based on waveform-resolved beta burst rate offer better decoding performance and improve the decoding speed versus accuracy trade-off when compared to standard band-limited power-based classification features.

## Materials and Methods

### Datasets

We analyzed the recordings of six open EEG MI benchmark datasets available through the MOABB (Aristimunha et al., 2023; Jayaram and Barachant, 2018) project: BNCI 2014-001 (Tangermann et al., 2012), BNCI 2014-004 (Leeb et al., 2007), Cho 2017 (Cho et al., 2017), MunichMI (Grosse-Wentrup 2009) (Grosse-Wentrup et al., 2009), Weibo 2014 (Yi et al., 2014) and Zhou 2016 (Zhou et al., 2016), and the recordings of a composite open EEG dataset that became recently available (Dreyer et al., 2023) referred to hereafter as Dreyer 2023. All datasets comprise a number of subjects with recordings corresponding to multiple trials of two or more randomly chosen, sustained kinesthetic MI commands, each performed following the appearance of a visual cue on a screen (Table 1). For our analysis we only considered trials corresponding to the “left hand” or “right hand” classes.

**Table 1.**
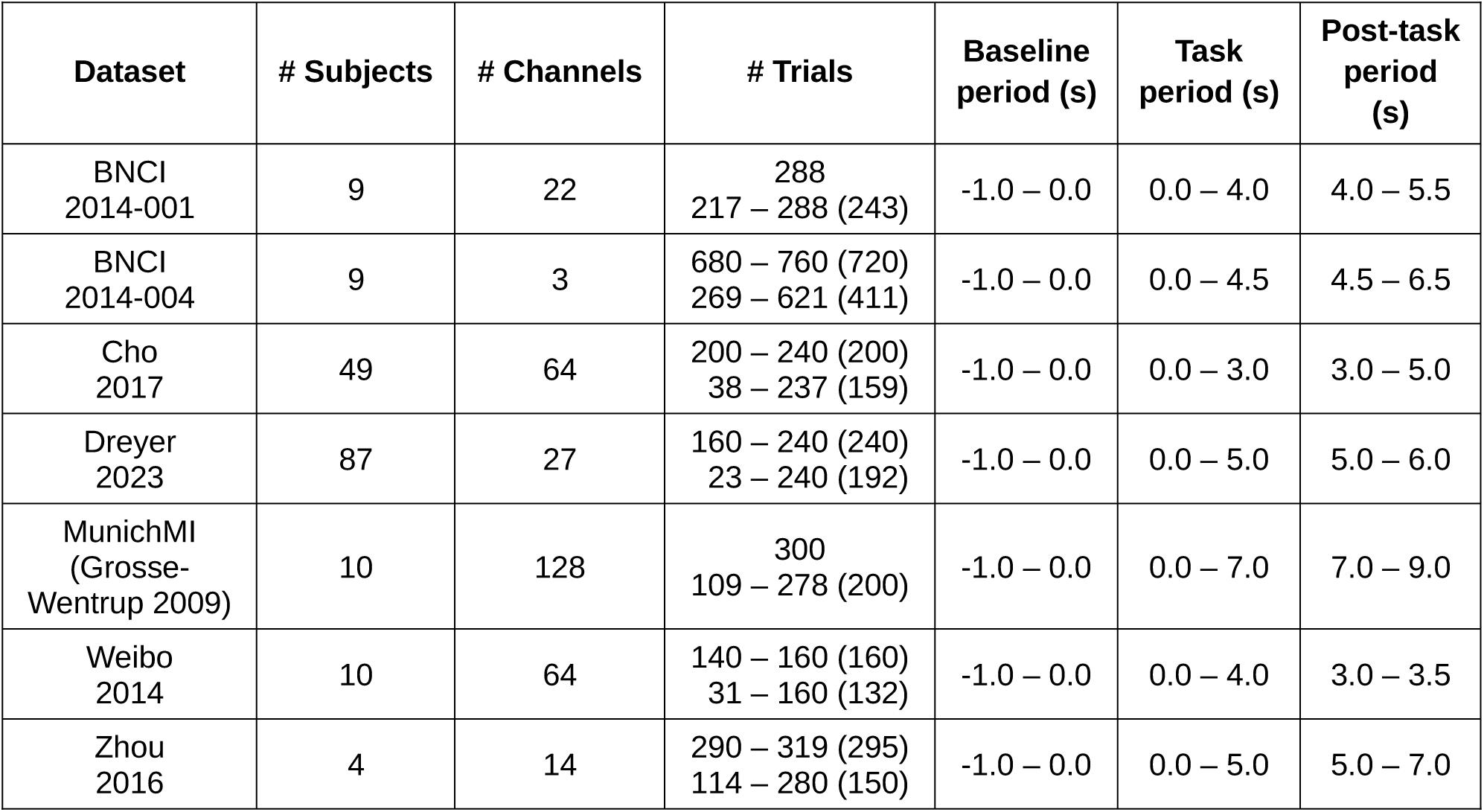
Dataset attributes. The lines in the fourth column indicate the original number of trials per subject (or the range in case this number was different between subjects), and the range of remaining trials across all subjects following trial rejection. Numbers in parentheses indicate the median number of trials.

### Pre-processing

The epoched recordings of each subject were loaded using the MOABB python package (v0.4.6, class LeftRightImagery; parameters: t_min_ and t_max_ as in indicated in Table 1), and were filtered with a low pass cutoff of 120 Hz (parameters: f_min_ = 0, f_max_ = 120; default MNE (Gramfort et al., 2013) zero-phase FIR filter designed with the windowed approach and transition bandwidth of 25% of the low pass frequency). Because the sampling frequency of the Weibo 2014 recordings is 200 Hz, we set the low pass cutoff to 95 Hz for this dataset. Finally, we used the autoreject python package (Jas et al., 2017) (v0.4.0, function get_rejection_threshold, default para-meters) in order to remove noisy trials.

### Burst detection and kernel selection

In order to select kernels for convolving the data from all channels we first detected bursts from channels C3 and C4 (or equivalently channels 43 and 44 for the MunichMI (Grosse-Wentrup 2009) dataset). To do so, we applied the pre-processing steps described above within a dataset-specific cluster of channels above the sensorimotor cortex (Papadopoulos et al., 2024). We applied a time-frequency (TF) decomposition in the 1 – 43Hz range on each selected channel separately, using the superlets algorithm (Moca et al., 2021) (parameters: o_min_ = 1, o_max_ = 40, c = 4) with a frequency resolution of 0.5 Hz. We noted high-power artifacts at approximately 25 – 30 Hz when inspecting the TF of the Cho 2017, Dreyer 2023 and MunichMI (Grosse-Wentrup 2009) datasets. This noise interferes with the burst detection step, therefore, we included an extra pre-processing step, prior to trial rejection, based on a custom implementation of the ZapLine algorithm from the meegkit python package (de Cheveigné, 2020) (v0.1.3, dss_line function) to remove these artifacts. Then, we detected bursts within the beta frequency range (15 – 30 Hz) from each TF matrix channel and used their temporal location to extract their waveforms from the raw time series within a fixed time-window of 260ms. For more details regarding the burst detection step we refer the readers to previous work from our group (Szul et al., 2023).

As the number of detected bursts per subject is large, we randomly sampled 10% of the trials of each participant per dataset and created a matrix that contained the waveforms of all detected bursts regardless of the trial class (“left” or “right” hand) for a given dataset. Due to the large number of subjects of the Dreyer 2023 dataset we restricted the random sample to 5% of each subject’s trials. Then, after robust scaling (scikit-learn package (Pedregosa et al., 2011), v1.0.2), we reduced the time dimension of the waveforms using principal component analysis (PCA) (Shlens, 2014) (scikit-learn package, v1.0.2).

We used the PCA score of each waveform detected from electrodes C3 and C4 (or equivalently channels 43 and 44 for the MunichMI (Grosse-Wentrup 2009) dataset), which is a metric of the difference between any waveform and the average shape of all bursts contained in the matrix provided as input to PCA. We defined an index of lateralized modulation of the average-per-axis PCA score *I_m_* :

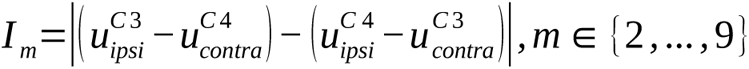

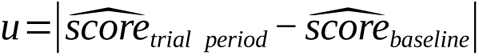

where ipsi (contra) refers to bursts recorded from channels C3 / C4 during a left / right (right / left) hand MI (Figure 1 a).

**Figure 1:**
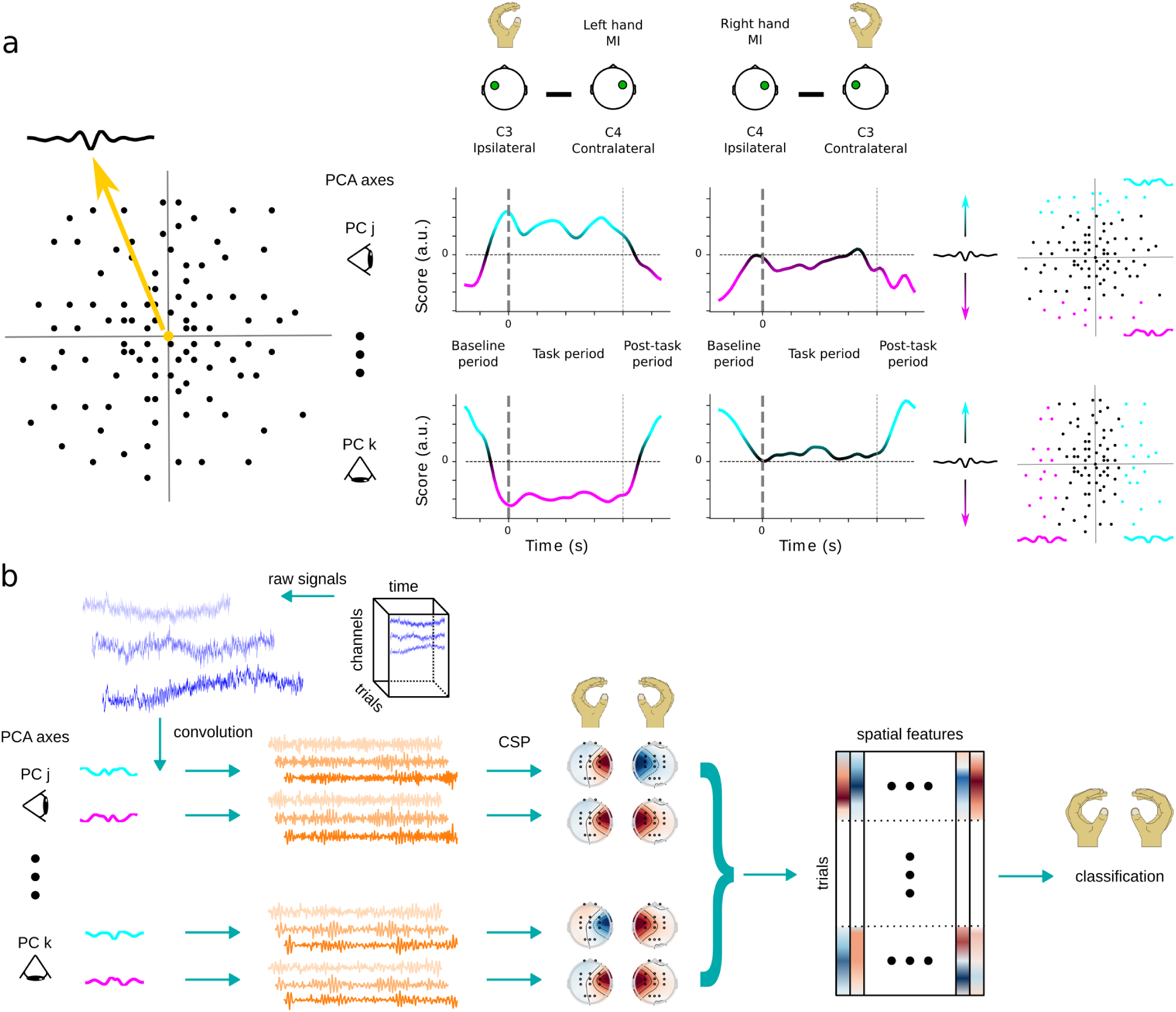
Illustration of methodology for computing classification features based on the convolution of raw signals with beta burst waveform kernels. **(a)** After randomly sampling the recording trials of all subjects within any dataset the beta bursts waveforms are analyzed using PCA. This constructs a high-dimensional space whose origin corresponds to the shape of the average waveform or equivalently a score equal to 0, and each axis defines a different axis of waveform variation. By only considering the beta bursts of channels C3 and C4 that occur at any point in time, the lateralization modulation index *Im* dynamically identifies the expected deviation of the average waveform shape from the overall average shape for each PCA axis. The axes that maximize *Im* are identified, the bursts are projected on these axes, split in groups of similarly shaped waveforms and the average waveform shapes of the two extrema are computed. **(b)** The raw signals of all recording channels of each dataset are independently convolved with each selected waveform from **a** resulting in distinctly temporally filtered copies of the signals. Each copy is then spatially filtered using the CSP algorithm, and finally all spatial features are concatenated in a single matrix that is provided as input to the classifier.

This index measures the inter-hemispheric difference of the average waveform shape between the baseline and trial periods. Its values span the range [0, ∞) and higher values indicate greater discrepancies between hemispheres and the two recording periods.

Based on observations from our previous study (Papadopoulos et al., 2024), we computed *I_m_* among components 2 to 9 in order to find three PCA axes that maximized this metric. We did not take into account the first component because it likely describes the temporal skew of the bursts (Papadopoulos et al., 2024; Szul et al., 2023). Finally, we divided the score range of each of the three selected axes in seven equally spaced groups, each group corresponding to a set of “similarly shaped” bursts. We kept the two groups per axis that lie further away from the origin (score equal to 0) and, by computing the Euclidean-average waveform of bursts within each group, we identified two kernels per axis. As such, we identified six kernels per dataset corresponding to burst waveforms whose rates were expected to be maximally modulated during the task, compared to baseline.

### Feature extraction

For each subject we applied the pre-processing, burst detection and kernel selection steps described above (pre-processing was applied to all available recording channels). Then, we convolved the EEG recordings with the corresponding kernels thus computing a proxy of the waveform-resolved burst rate per kernel. The temporally convolved, epoched data was then spatially filtered using the CSP algorithm (MNE package, v1.5.1, function CSP, parameters: n_components = 4, transform_into = “average_power”). Finally, we concatenated all 24 spatial features into a single vector for each trial (Figure 1 b).

To compare with, we also used standard approaches to compute spatial features of band-limited power modulations. After pre-processing, we independently filtered the epoched data in the mu (6 – 15 Hz), beta (15 – 30 Hz) or both the mu and beta (6 – 30 Hz) bands, using either a single filter or a filter bank approach. Then, the filtered data served as inputs to the CSP algorithm (using the already described parameters) resulting in four spatial features per filter. For the filter bank approach, we split either frequency range in non-overlapping filter banks with a frequency span of 3 Hz per filter. As such we defined three filters for the mu band (6 – 9 Hz, 9 – 12 Hz, 12 – 15 Hz), five filters for the beta band (15 – 18 Hz, 18 – 21 Hz, 21 – 24 Hz, 24 – 27 Hz, 27 – 30 Hz) and eight filters for the mu-beta band (6 – 9 Hz, 9 – 12 Hz, 12 – 15 Hz, 15 – 18 Hz, 18 – 21 Hz, 21 – 24 Hz, 24 – 27 Hz, 27 – 30 Hz). Then, we again used CSP and concatenated all spatial features of each filter bank, resulting in 12, 20 and 32 spatial features respectively per trial.

### Classification

We used a repeated (*n* = 10), 5-fold cross validation procedure to estimate the decoding score using linear discriminant analysis (LDA) (Tharwat et al., 2017; Vidaurre et al., 2011)(scikit-learn, v1.0.2) as a classifier. We adopted a time-resolved decoding paradigm, using both an incremental and a sliding time window. In the first case we started with a 100 ms time window and repeated the classification procedure by incrementing this window by 100 ms at a time. The baseline period was considered separately from the trial period. In the latter case we used 1 s long sliding time windows which moved in 50 ms increments. Decoding scores were based on the area under the curve (AUC) of the receiver operating characteristic (scikit-learn, v1.0.2). All numeric computations were based on the numpy python package (v1.21.6; (Harris et al., 2020)) and an environment running python (v3.10).

### Information transfer rate

Information transfer rate (Arslan and Sinha, 2024; Sadeghi and Maleki, 2019) was defined as:

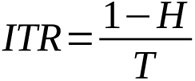

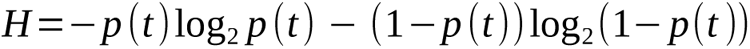

where the binary entropy function *H* depends on the average accuracy probability at any time window *p*(*t*), and *T* corresponds to the maximum recording time required by each time window in seconds. *T* was shifted such that all time values are positive, *i.e.* using the absolute time starting from the beginning of the baseline period (Table 1) when using a sliding window and when considering the baseline period using an incremental time window. The values of this metric span the range [0,10] when using an incremental window decoding approach and [0,20] when using a sliding window. Large values indicate a better trade-off between decoding accuracy and decoding speed.

### Statistical analysis

On the dataset level, we performed pairwise comparisons of the across-subject average decoding score corresponding to the beta burst convolution spatial features and each of the spatial features of the band-limited power modulations. These comparisons were based on threshold-free cluster-based permutation (*n* = 2^13^) tests (MNE package, v1.5.1, function permutation_cluster_test, parameters: threshold = dict(start = 0, step = 0.2), tail = 1) that were subsequently thresholded at significance level of a = 0.05 for visualization purposes.

To estimate, on the population level, any statistical differences between the maximum classification scores obtained using different feature extraction pipelines during the trial period, we compared the scores of the beta burst convolution pipeline against those based on classical filtering pipelines. We used a linear mixed model with across-trials average classification score as the dependent variable setting the number of trials as prior weights, the type of classification feature as a fixed effect, and subject nested within dataset as random intercepts. We implemented similar models to compare the time required to achieve the maximum classification score per feature extraction pipeline, and also the maximum ITR and time needed to reach it. In the latter two cases we first transformed the values to logarithmic scale in order to ensure normality of the residuals. Statistical analyses were conducted using R (v4.1.2) and lme4 (v1.1-31; (Bates et al., 2015)). Fixed effects were assessed using type II Wald X 2 tests using car (v3.1-1; (Fox and Weisberg, 2019)). Pairwise Tukey-corrected follow-up tests were carried out using estimated marginal means from the emmeans package (v.1,8,7; (Lenth, 2023)).

## Results

In summary, we have employed seven freely available datasets of EEG recordings from subjects performing left and right hand MI. Within each dataset, we detected beta bursts for each subject within electrode clusters over left and right sensorimotor cortex and then randomly sampled 10% of the trials containing these bursts (the sample size was limited to 5% for the Dreyer 2023 dataset). We applied PCA to the matrix of all burst waveforms and defined a modulation index *I_m_* in order to find burst waveform shapes whose lateralized rates were expected to be maximally modulated between the baseline and trial periods. These waveforms were then employed as kernels for convolving the EEG data in time domain before applying spatial filtering with CSP. Finally all spatial features were combined and served as input for LDA, in order to classify “left” versus “right” hand MI. We also performed classification by applying standard temporal filtering techniques before applying spatial filtering with CSP, using either a single filter or a filter bank in the mu (6 – 15 Hz), beta (15 – 30 Hz) and mu-beta (6 – 30 Hz) frequency bands. We estimated the time-resolved decoding score per subject of each dataset for each classification feature employing both an incremental and a sliding decoding window (see Materials and Methods for details).

Across all datasets, the average decoding accuracy obtained using the proposed methodology based on beta bursts outperformed the results based on standard beta band filtering irrespective of the filtering (single filter or filter bank) or the windowing (incremental or sliding) technique during most of the recording time (Figure 2 a, Sup. Figure 1 a). Within each dataset, the across-subjects average score obtained by beta bursts was higher than that of any beta band filtering technique, usually shortly after the beginning of the trial or towards its end (Figure 2 b-h, Sup. Figure 1 b-h). Exceptions to this finding when using an incremental window were the BNCI 2014-001 dataset, for which all features produced equivalent results (Figure 2 c, Sup. Figure 1 c), and the Weibo 2014 dataset, for which mu-beta filtering outperformed beta bursts (Figure 2 g, Sup. Figure 1 g). Threshold-free cluster-based permutation tests (see Materials and Methods) revealed a significant cluster of increased accuracy for beta bursts compared to either beta filtering technique during most of the recording time or following the trial onset for each windowing (incremental or sliding) technique respectively (Figure 2 a, Sup. Figure 1 a). Within each dataset, we found clusters similar to those of the population average for the Dreyer 2023 dataset (Figure 2 e, Sup. Figure 1 e). The across-subjects average score obtained by beta bursts was higher than that of any beta band filtering technique shortly after the beginning of the trial for the BNCI 2014-001 and Cho 2017 datasets (Figure 2 b, d, Sup. Figure 1 b, d). For the rest of the datasets (BNCI 2014-004, MunichMI (Grosse-Wentrup 2009), Weibo 2014, Zhou 2016) no differences were observed among the beta bursts and either beta band filtering method. Overall, in terms of classification accuracy, we observed an improvement with beta bursts over beta power on the population level and in 5/7 (4/7) datasets, with clusters of statistically significant differences arising on the population level and in 3/7 (2/7) datasets when using an incremental (sliding) window.

**Figure 2:**
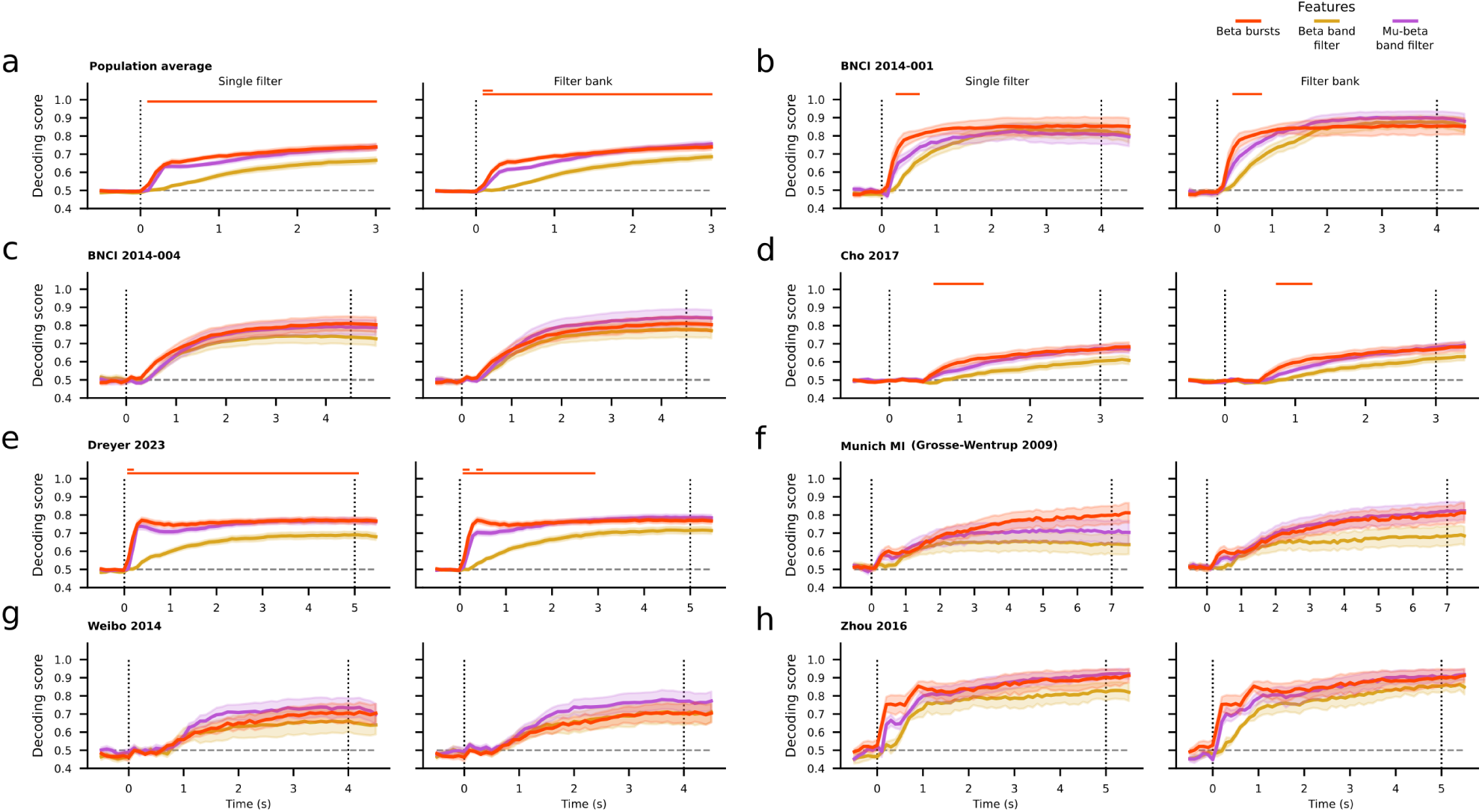
**(a)** Population average time-resolved decoding score and standard error for the beta burst convolution (red), beta band (yellow) and mu-beta band (purple) filtering pipelines using an incremental window. Due to the different duration of the task per dataset we restricted the time to the minimum trial period corresponding to 3 seconds. **(b – h)** Average, time-resolved decoding score and standard error per dataset of the same features using an incremental window. For each panel, the left subplot depicts the decoding results obtained using a single filter, while the right subplot depicts the results based on a filter bank technique. The beta burst results are the same for the pair of each panel. The horizontal dashed line corresponds to the expected chance level. Vertical dotted lines represent the onset and end of the trial period of each dataset. Horizontal lines on the top of each subplot show the results of pair-wise permutation cluster tests between the beta bursts and either filtering technique, with correction for multiple comparisons at significance level of 0.05. The color of the lines indicate which feature produces, on average, better results at any time point. A lack of color indicates no statistically significant differences between the compared features.

We did not observe such clear differences when comparing the beta bursts and mu-beta filtering decoding scores. On the population level, when using an incremental time window, average beta burst convolution results outperformed the single filter and filter bank techniques early after the beginning of a trial (Figure 2 a). On the dataset level, this was true for the BNCI 2014-001 and Dreyer datasets (Figure 2 b, e). For the rest of the datasets, differences varied depending on the filtering technique. Notably, in the mu-beta band, both filtering techniques produced higher decoding scores than beta bursts for the Weibo 2014 dataset (Figure 2 g). A similar pattern was also observed when using a sliding window (Sup. Figure 1 a-h). The permutation tests between the beta burst features and the mu-beta filtering techniques revealed only small clusters on the population level, as well for the Dreyer 2023 dataset. We found that beta bursts improve classification scores over the mu-band filtering techniques on the population level and 4/7 (2/7) datasets when using an incremental (sliding) window, with small clusters of statistically significant differences on the population level and one dataset only when using an incremental window.

Finally, comparisons between beta burst convolution and mu filtering results showed an improvement when using a sliding window. Clusters of statistically significant differences revealed that beta bursts are better than single-filter mu band power on the population level and 2/7 datasets (Sup. Figure 2 a-c). No differences were found on the population level when comparing beta bursts to the filter bank technique, but on the dataset level the beta bursts score was better for the BNCI 2014-001 dataset (Sup. Figure 2 b) and conversely worse for the Weibo 2014 dataset (Sup. Figure 2 g) based on cluster permutation tests. Comparisons of results when using a sliding window approach did not reveal any differences (Sup. Figure 3).

For all datasets we computed the information transfer rate (ITR) in order to quantify the difference between all classification features in terms of the decoding speed-accuracy trade-off (see Materials and Methods). On the population level, beta bursts provided a higher ITR than either beta band filtering technique across the whole trial (Figure 3 a). On the dataset level, the same pattern was observed for the Dreyer 2023 dataset (Figure 3 e). Moreover, beta bursts ITR was higher than any beta band filtering early after trial onset for the BNCI 2014-001, Munich MI (Grosse-Wentrup 2009) and Zhou 2016 datasets, and later for the Cho 2017 dataset. No differences between the features were observed in the case of BNCI 2014-004 dataset, whereas beta bursts resulted in the lowest ITR for the Weibo 2014 dataset (Figure 3 b-h). Permutation cluster tests revealed a significant difference between beta bursts and either filtering technique in the beta band on the population level, and for most of the time for the Dreyer 2023 dataset. A cluster after the trial onset was found for the BNCI 2014-001 dataset, and no clusters were found for the rest of the datasets. When using a sliding window approach, beta bursts provided a higher ITR than either beta band filtering technique on the population level early after the beginning of the trial and towards its end highlighted by the presence of a cluster (Sup. Figure 4 a). A significant difference was also found for the Dreyer 2023 dataset after the trial onset (Sup. Figure 4 e). For datasets BNCI 2014-001, Cho 2017, Dreyer 2023 and Zhou 2016 we observed a higher ITR for beta bursts compared to beta band filtering mainly after the trial onset, whereas for the rest of the datasets either feature resulted in equivalent ITRs (Sup. Figure 4 b-h). No significant clusters were found for any of these datasets. In summary, irrespective of the windowing method beta bursts resulted in higher ITR than beta power on the population level and for 5/7 datasets with clusters arising on the population level and 2/7 datasets.

**Figure 3:**
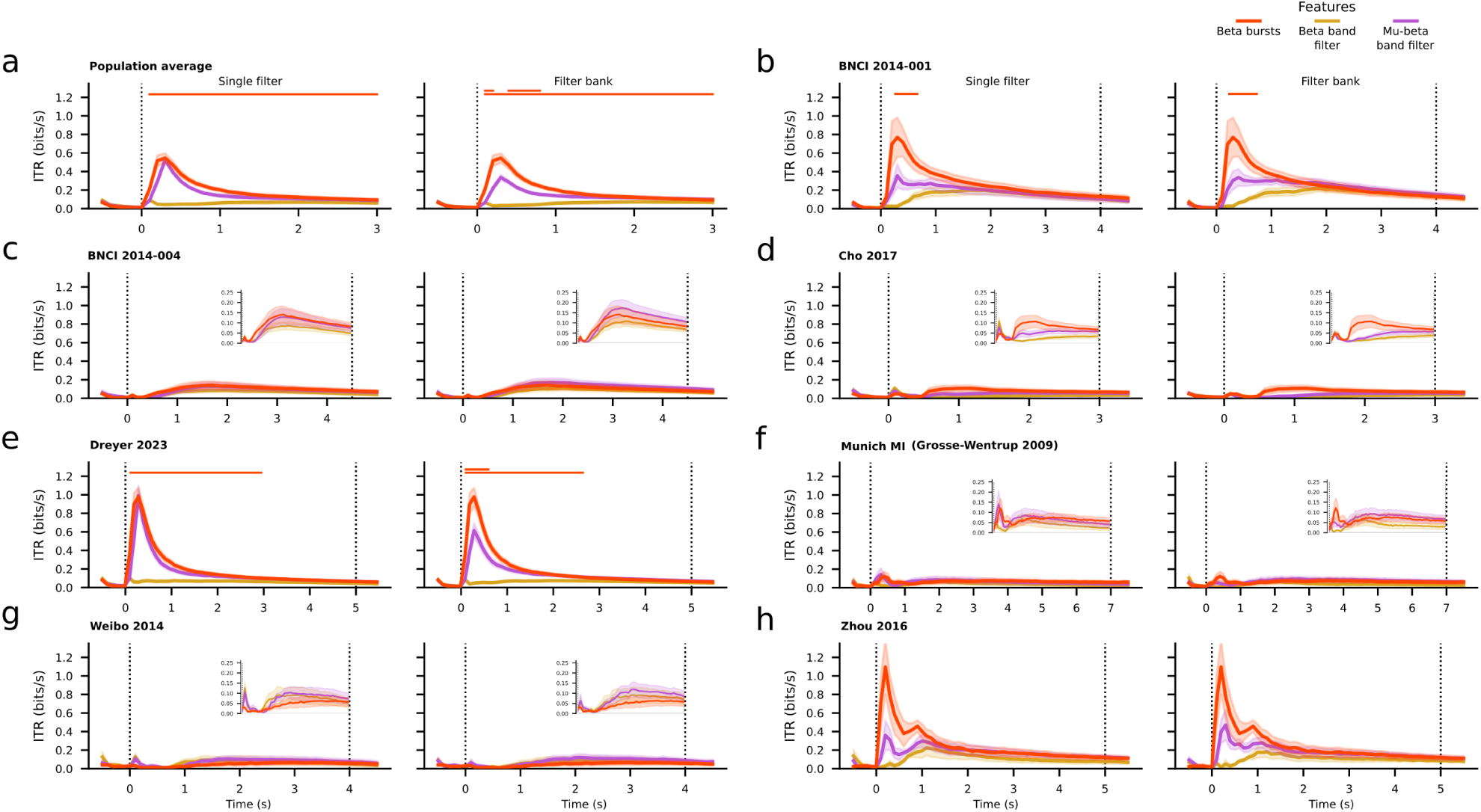
**(a)** Population average time-resolved information transfer rate (ITR) and standard error for the beta burst convolution (red), beta band (yellow) and mu-beta band (purple) filtering pipelines using an incremental window. Due to the different duration of the task per dataset we restricted the time to the minimum trial period corresponding to 3 seconds. **(b – h)** Average, time-resolved information transfer rate (ITR) and standard error per dataset of the same features using an incremental window. For each panel, the left subplot depicts the ITR results obtained using a single filter, while the right subplot depicts the results based on a filter bank technique. The beta burst results are the same for the pair of each panel. Vertical dotted lines represent the onset and end of the trial period of each dataset. Horizontal lines on the top of each subplot show the results of pair-wise permutation cluster tests between the beta bursts and either filtering technique, with correction for multiple comparisons at significance level of 0.05. The color of the lines indicate which feature produces, on average, better results at any time point. A lack of color indicates no statistically significant differences between the compared features. Insets provide a zoomed-in view of the values within the corresponding panels.

Regarding the comparison between beta bursts and the mu-beta filtering techniques, ITR was higher for the former on the population level as well as datasets BNCI 2014-001, Cho 2017, Dreyer 2023, Munich MI (Grosse-Wentrup 2009) and Zhou 2016 shortly after the trial onset, especially when adopting the filter bank method. No differences were observed for the BNCI 2014-004 dataset. The mu-band filter bank technique produced higher ITR than beta bursts period in the case of the Weibo 2014 dataset (Figure 3 a-h). The results were similar when using a sliding window (Sup. Figure 4 a-h). On the population level, cluster-based permutations tests revealed a significant difference between the beta bursts features and the filter bank in the mu-beta band using either an incremental or a sliding window approach, but no significant clusters when comparing the beta bursts to the single filter in the mu-beta band (Figure 3 a, Sup. Figure 4 a). Similar clusters of significant differences between the features were found only for the Dreyer 2023 dataset with an incremental window (Figure 3 e, Sup. Figure 4 e). Overall, beta bursts yielded higher ITR than mu-beta power on the population level and 4/7 datasets regardless of the windowing technique. Permutation cluster tests revealed improvements attributable to beta bursts compared to the filter bank technique on the population level, and for one dataset when using an incremental time window.

We used linear mixed models to quantify the differences in maximum decoding score, latency to achieve the maximum score, maximum ITR and latency to the maximum ITR per feature (see Materials and Methods) using both the incremental and sliding windows (Figure 4, Sup. Figure 5).

**Figure 4:**
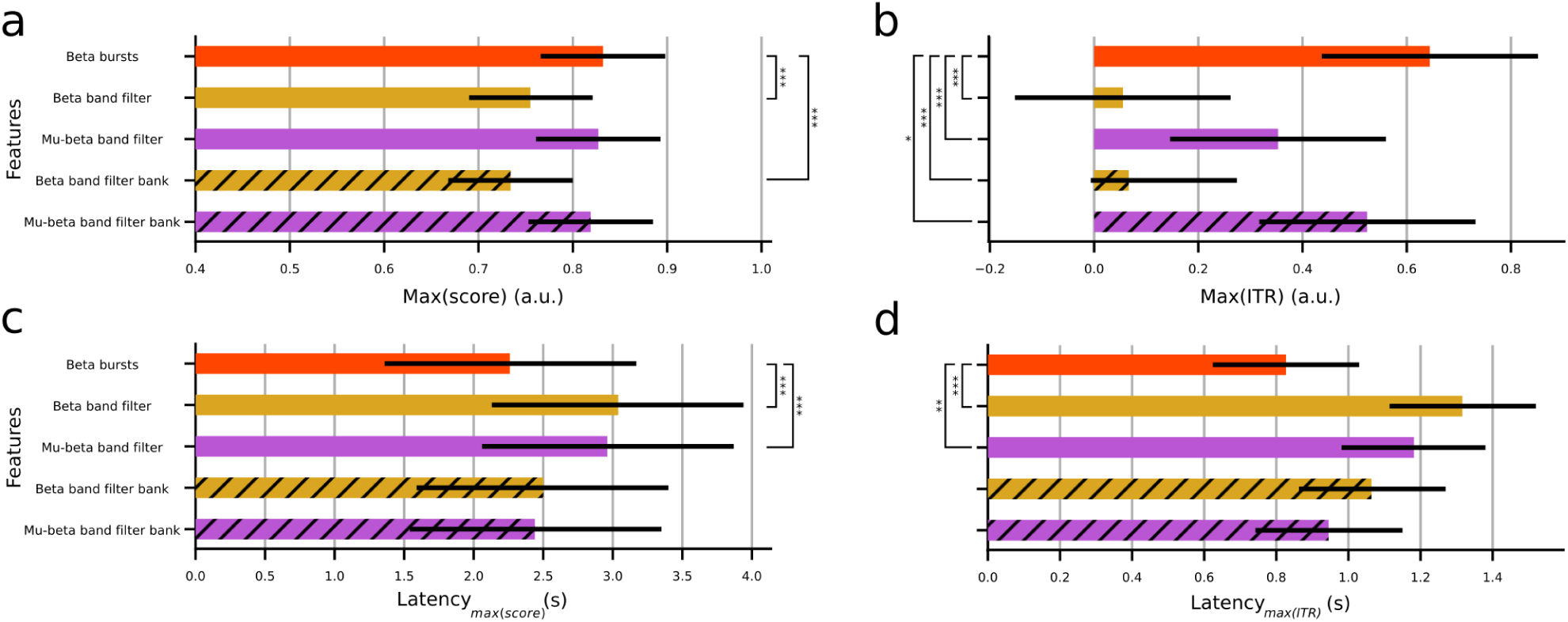
Population-level statistical analysis based on linear mixed models when using an incremental window per feature. **(a)** Average maximum decoding score. **(b)** Average maximum ITR. **(c)** Average latency to reach the maximum decoding score. **(d)** Average latency to reach the maximum ITR. Error bars show 95% confidence intervals. Hatches indicate the use of a filter bank technique. Asterisks indicate statistically significant differences among pairwise comparisons of the beta bursts and the rest of the features (* : p < 0.05. ** : p < 0.01, *** : p < 0.001). A lack of asterisks implies no statistically significant differences. Note that the log transform of the the maximum ITR and latency to maximum ITR were used for the statistical analysis (see Materials and Methods), but panels **b** and **d** depict results before applying the transformation for ease of comparisons with panels **a** and **c** respectively.

On the population level, the maximum classification scores for the beta bursts technique, filtering in the beta band using a single filter, filtering in the mu band using a single filter, and filter bank in the beta or mu and beta bands using an incremental time window were 0.832 ± 0.066, 0.755 ± 0.065, 0.827 ± 0.066, 0.734 ± 0.066 and 0.819 ± 0.066 (Figure 4 a). The time required to reach each of these decoding scores (latency) was 2.26 ± 0.91, 3.04 ± 0.90, 2.96 ± 0.90, 2.50 ± 0.90 and 2.44 ± 0.91 seconds (Figure 4 c), respectively. The maxima of the ITR before applying the logarithmic transformation (see Materials and Methods) per feature were 0.6443 ± 0.207, 0.0554 ± 0.207, 0.3531 ± 0.0206, 0.0667 ± 0.206, 0.5241 ± 0.207 (Figure 4 b). The corresponding average latencies before the logarithmic transformation were 0.827 ± 0.203, 1.316 ± 0.204, 1.182 ± 0.203, 1.064 ± 0.201, 0.945 ± 0.203 seconds (Figure 4 d). Regarding the analysis based on a sliding window, the maximum classification scores were 0.816 ± 0.050, 0.753 ± 0.050, 0.810 ± 0.050, 0.753 ± 0.050 and 0.818 ± 0.050 respectively (Sup. Figure 5 a). The latencies were 2.03 ± 0.23, 2.45 ± 0.22, 2.19 ± 0.23, 2.49 ± 0.24 and 2.22 ± 0.23 seconds (Sup. Figure 5 c). Before applying the logarithmic transformation, the ITR maxima were 0.694 ± 0.180, 0.170 ± 0.178, 0.413 ± 0.177, 0.206 ± 0.182, 0.533 ± 0.179 (Sup. Figure 5 b), and the corresponding latencies 0.575 ± 0.162, 0.840 ± 0.161, 0.796 ± 0.160, 0.743 ± 0.161, 0.701 ± 0.162 seconds (Sup. Figure 5 d).

Across all datasets the maximum classification accuracy of the beta bursts technique when using an incremental window was significantly higher than that of the beta band single filter and filter bank pipelines (*X^2^(4)* = 326.81, *t(24587436)* = 10.697, *p* < 0.001 and *t(24548972) = 13.705*, *p* < 0.001 respectively), but did not differ significantly from either technique exploiting both the mu and beta bands (*t(24904394)* = 0.637, *p* = 0.9691 and *t(24049399)* = 1.777, *p* = 0.3873). The latency to achieve the maximum score was significantly lower for the beta burst convolution pipeline compared to both the single filtering techniques in the beta and mu-beta bands (*X^2^(4)* = 62.508, *t(24698263)* = -6.361, *p* < 0.001 and *t(25032435)* = -5.769, *p* < 0.001), but did not differ significantly compared to the corresponding filter bank techniques (*t(24649522)* = -1.926, *p* = 0.3034 and *t(24151454)* = -1.488, *p* = 0.5703). The logarithmic transform of the maximum ITR for the beta bursts technique was significantly higher than the single filtering and filter bank techniques in both the beta and mu-beta bands (*X^2^(4)* = 309.58, *t(24722157)* = 14.0967, *p* < 0.001, *t(24671284)* = 12.904, *p* < 0.001 and *t(25060057)* = 5.127, *p* < 0.001, *t(24173608)* = 2.909, *p* = 0.0298). The logarithmic transform of the latency to achieve maximum ITR was significantly lower for the beta bursts technique compared to either single filtering pipeline in the beta and mu-beta bands (*X^2^(4)* = 40.75, *t(24777354)* = -5.863, *p* < 0.001 and *t(25130735)* = -3.885, *p* = 0.001), but did not significantly differ compared to the filter bank method in either band (*t(24721433)* = -2.071, *p* = 0.2329 and *t(24227058)* = -1.545, *p* = 0.5329).

When using a sliding window, across all datasets the maximum classification accuracy of the beta bursts technique was significantly higher than that of the beta band single filter and filter bank pipelines (*X^2^(4)* = 237.95, *t(24582761)* = 10.072, *p* < 0.001 and *t(24544732) = 10.055*, *p* < 0.001 respectively), but did not differ significantly from either technique exploiting both the mu and beta bands (*t(24898964)* = 0.956, *p* = 0.8748 and *t(24045211)* = -0.418, *p* = 0.9936). Similarly, the latency to achieve the maximum score was significantly lower for the beta burst convolution pipeline compared to either filtering technique in the beta band (*X^2^(4)* = 25.552, *t(24736767)* = - 3.890, *p* < 0.001 and *t(24684274)* = -4.281, *p* < 0.001), but did not differ significantly compared to the mu-beta band (*t(25086359)* = -1.467, *p* = 0.5840 and *t(24189739)* = -1.762, *p* = 0.3961). The logarithmic transform of the maximum ITR for the beta bursts technique was significantly higher than both filtering techniques in the beta band and the single filtering in the mu-beta band (*X^2^(4)* = 162.26, *t(24732475)* = 10.236, *p* < 0.001, *t(24680636)* = 9.172, *p* < 0.001 and *t(25072582)* = 4.499, *p* < 0.001) but did not significantly differ from the mu-beta filter bank (*t(24183763)* = 1.665, *p* = 0.4559). The logarithmic transform of the latency to achieve maximum ITR was significantly lower for the beta bursts technique compared to either single filtering pipeline in the beta and mu-beta bands (*X^2^(4)* = 13.151, *t(24808114)* = -3.069, *p* = 0.0183 and *t(25164334)* = -3.029, *p* = 0.0207), but did not significantly differ compared to the filter bank method in either band (*t(24749694)* = -1.796, *p* = 0.3756 and *t(24255336)* = -1.296, *p* = 0.6937).

## Discussion

Standard techniques for analyzing meso-and macro-scale neural signals recorded during the execution or imagination of movements typically rely on signal power metrics assuming that relevant changes in brain signals are reflected in amplitude modulation (Alayrangues et al., 2019; Kilavik et al., 2013; Pfurtscheller, 1981; Pfurtscheller et al., 1997, 1996; Pfurtscheller and Lopes da Silva, 1999). However, there has recently been a considerable paradigm shift towards considering transient signal features on the single trial level (Chen et al., 2021; Coleman et al., 2024; Jones, 2016; Little et al., 2019; Lundqvist et al., 2024, 2016; Rayson et al., 2023; Shin et al., 2017; Szul et al., 2023; Torrecillos et al., 2018; Vigué-Guix and Soto-Faraco, 2022; Wessel, 2020). Therefore, considering that computational models describing the neuronal generators of specific burst waveform shapes (Bonaiuto et al., 2021; Sherman et al., 2016; Szul et al., 2023) offer an improved theoretical interpretability of the observed signal modulations, applications leveraging such signal characteristics, like beta bursts, could potentially benefit from incorporating recent neuroscience findings (Papadopoulos et al., 2022).

In this work we analyzed the activity of seven open EEG MI datasets and focused on developing a streamlined process for incorporating beta bursts into a BCI pipeline. We defined a modulation index *I_m_* which allowed us to identify beta burst waveforms whose rate is expected to be modulated due to task demands. This was based on the assumption that maximal modulations of the average waveform shape along specific PCA components are the result of a net imbalance of the rates of bursts with different shapes driven by task demands. PCA constitutes a mathematically tractable and interpretable way of analyzing the diversity of beta burst waveform shapes, but other supervised dimensionality reduction algorithms could better disentangle the waveforms by imposing different constraints, e.g. taking into account trial labels.

We used these waveforms as kernels to convolve the raw signals in the time domain. We chose to keep the number of kernels and the method for extracting them fixed based on insights from a previous study (Papadopoulos et al., 2024), but future work could employ a formal hyper-parameter search so as to maximize classification accuracy. Similarly, we did not check for any redundancy in the convolved signals due to selection of similarly shaped kernels, a point which could be addressed by another dimensionality reduction algorithm. The implementation of the convolution is virtually as computationally efficient as any filtering technique. However, we note that the proposed methodology assumes the existence of data that can be analyzed offline in order to first find the relevant beta burst waveforms to use as kernels. Moreover, this data needs to be clean of artifacts as the beta burst detection algorithm may fail to detect transient activity in the presence of high-power oscillations, instead detecting only the latter. For this reason, we believe that the superlets algorithm is the only time-frequency decomposition technique providing adequate time and frequency resolution in order to determine if the data is clean and to extract burst waveforms.

This data-driven, neurophysiology-informed filtering of the signals resulted in a proxy of waveform-resolved burst rate per kernel. We also performed a standard filtering in the mu (6 – 15 Hz) band, beta (15 – 30 Hz) band, or a wider frequency range encompassing both the mu and beta (6 – 30 Hz) bands. Finally, we used CSP to extract spatial features for classification. We showed that classification scores can be improved compared to a standard power-based analysis of the beta band activity, requiring briefer recordings to do so and that, without explicitly considering the mu (6 – 15 Hz) band activity in our beta bursts pipeline, the corresponding classification scores are equivalent to scores of state-of-the-art approaches again needing shorter recordings. Further, the filter bank-based analysis allowed us to verify that the decoding improvements where not simply the result of an increase in number of spatial features used for classification. Instead, beta burst waveforms are more informative of the underlying MI task on the population level than beta band power, and equally informative to standard power-based techniques that take into account mu activity modulations. A possible explanation for this is that the beta burst kernels also capture slower modulations of the underlying activity. Future work could explore the incorporation of novel mu band features or a combination of mu power and beta burst features.

By employing both an incremental and a sliding window strategy for classification we can speculate on the observed differences of decoding scores across classification features and datasets. First, it appears that MI does not begin right after the go cue in all datasets but can be delayed possibly due to differences in the task design or the instructions given to the participants. Second, relatively sustained decoding performances when using a sliding window were translated into slow increases of those performances when using an incremental window. In contrast, a drop in performance when using a sliding window was reflected in a plateau when using an incremental window. Taken together these observations imply that finding an optimal decoding time window, especially for online paradigms, is not trivial and depends not only on the selected classification features or algorithm but also on experimental design variables.

In order to assess the trade-off between decoding accuracy and speed, we used the classification scores obtained by implementing each pipeline and computed the corresponding ITR. We showed a statistically significant increase of the maximum ITR achieved using the beta burst kernel filtering compared to any other method, and a statistically significant decrease of the time needed to achieve this value compared to single filters. These results suggest that beta bursts could be particularly relevant for BCI applications that aim to minimize the recording time required before issuing a command like in the case of real-time decoding of a switch control, although the non-structured nature of such applications still poses a considerable challenge (Carrara and Papadopoulo, 2024).

This study focused on incorporating recent neurophysiology insights, specifically task dependent waveform-specific modulation of beta burst rates, in a pipeline for decoding EEG signals during imagined movements. We proposed a simple and computationally efficient algorithm that leverages beta burst waveforms and transforms brain recordings in a way that is compatible with other widely adopted algorithms. We demonstrated that classification results based on beta bursts are superior to results based beta-band power alone, and are on-par with power-based results that take into account the mu band. By computing the information transfer rate we showed that, often, features based on beta bursts significantly improve the decoding speed-accuracy trade-off. We also verified this finding using a sliding window decoding technique, a fact which further suggests the feasibility for online decoding with this approach. Taking everything into account, we believe that these findings can serve as an important step in the direction of improving online BCI decoding paradigms.

## Data and Code Availability

All data are freely available via the <u>MOABB project</u> and the open access repository Zenodo at https://zenodo.org/records/8089820. All scripts necessary for reproducing the results of this article are available at the following public repository: https://gitlab.com/sotpapad/bebopbci/-/tree/preprint_version_202407.

## Author Contributions

S P, J B and J M conceptualized the manuscript. S P drafted the manuscript and performed the analysis. All authors contributed to manuscript revision, read, and approved the submitted version.

## Funding

S P, L D, M C, J B and J M are supported by the French National Research Agency (ANR) project HiFi (2020–2024, ANR-20-CE17-0023). M S and J B are supported by the European Research Council (ERC) under the European Union’s Horizon 2020 Research and Innovation Programme (ERC consolidator Grant 864550 to J B). This work was performed within the framework of the LABEX CORTEX (ANR-11LABX-0042) of Université de Lyon, within the program ‘Investissements d’Avenir’ (decision n◦ 2019-ANR-LABX-02) operated by the French National Research Agency (ANR).

## Declaration of Competing Interests

All authors declare no competing interests.

---

**Sup. Figure 1:**
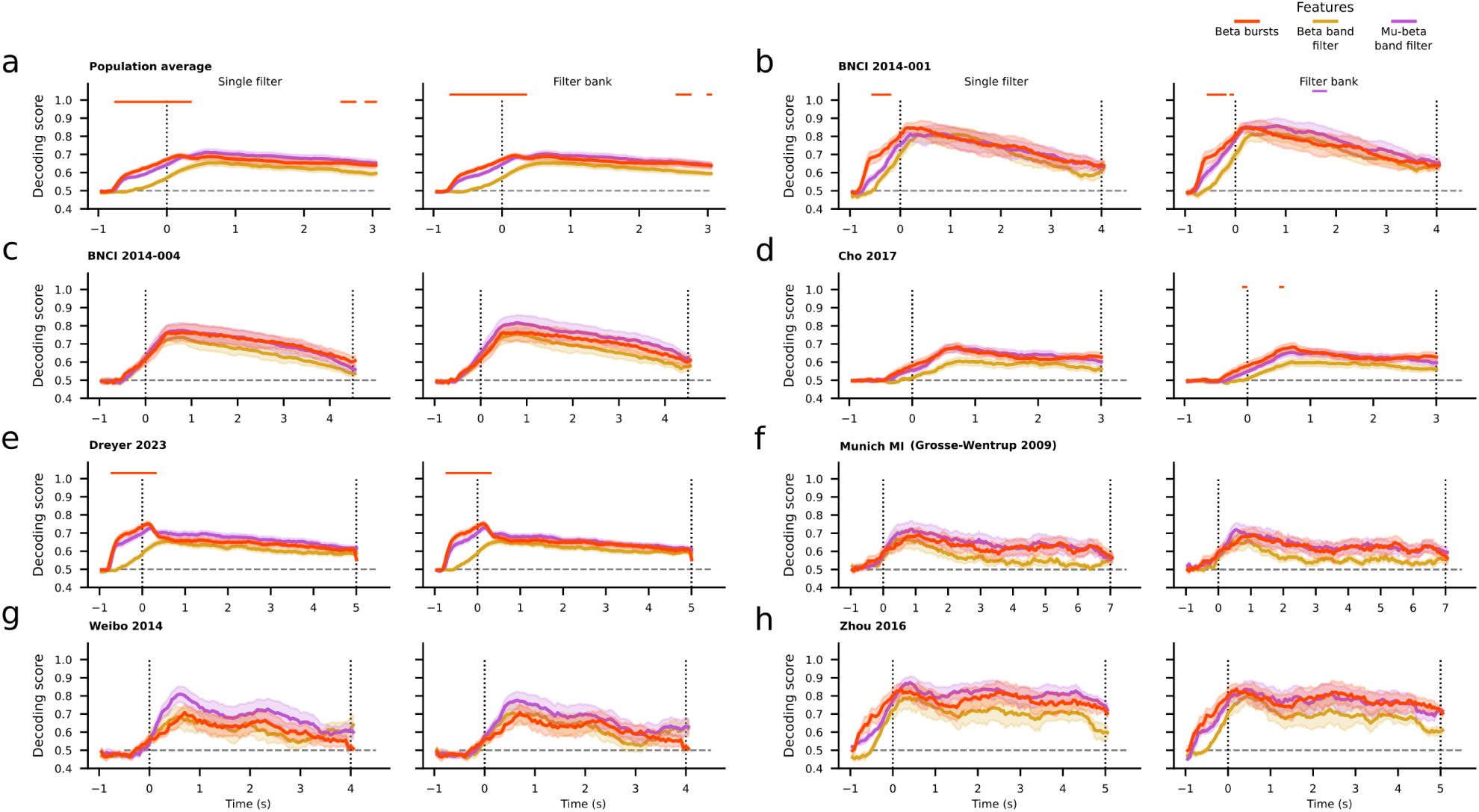
**(a)** Population average time-resolved decoding score and standard error for the beta burst convolution (red), beta band (yellow) and mu-beta band (purple) filtering pipelines using a sliding window. Due to the different duration of the task for each dataset we restricted the time to the minimum trial period corresponding to 3 seconds. **(b – h)** Average, time-resolved decoding score and standard error per dataset of the same features using a sliding window. For each panel, the left subplot depicts the decoding results obtained using a single filter, while the right subplot depicts the results based on a filter bank technique. The beta burst results are the same for the pair of each panel. The horizontal dashed line corresponds to the expected chance level. Vertical dotted lines represent the onset and end of the trial period of each dataset. Horizontal lines on the top of each subplot show the results of pair-wise permutation cluster tests between the beta bursts and either filtering technique, with correction for multiple comparisons at significance level of 0.05. The color of the lines indicate which feature produces, on average, better results at any time point. A lack of color indicates no statistically significant differences between the compared features.

**Sup. Figure 2:**
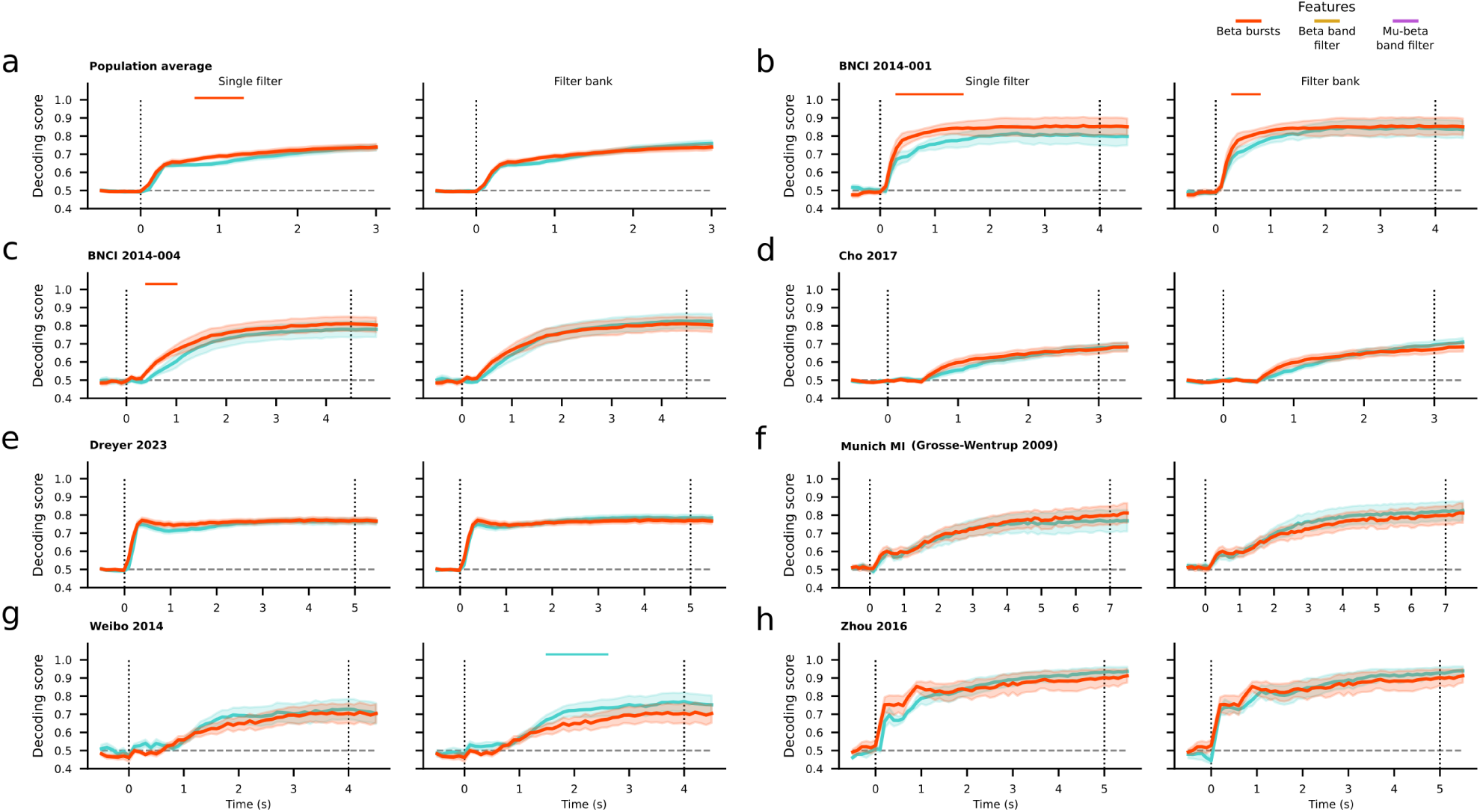
**(a)** Population average time-resolved decoding score and standard error for the beta burst convolution (red) and mu band (turquoise) filtering pipelines using an incremental window. Due to the different duration of the task for each dataset we restricted the time to the minimum trial period corresponding to 3 seconds. **(b – h)** Average, time-resolved decoding score and standard error per dataset of the same features using an incremental window. For each panel, the left subplot depicts the decoding results obtained using a single filter, while the right subplot depicts the results based on a filter bank technique. The beta burst results are the same for the pair of each panel. The horizontal dashed line corresponds to the expected chance level. Vertical dotted lines represent the onset and end of the trial period of each dataset. Horizontal lines on the top of each subplot show the results of pair-wise permutation cluster tests between the beta bursts and either filtering technique, with correction for multiple comparisons at significance level of 0.05. The color of the lines indicate which feature produces, on average, better results at any time point. A lack of color indicates no statistically significant differences between the compared features.

**Sup. Figure 3:**
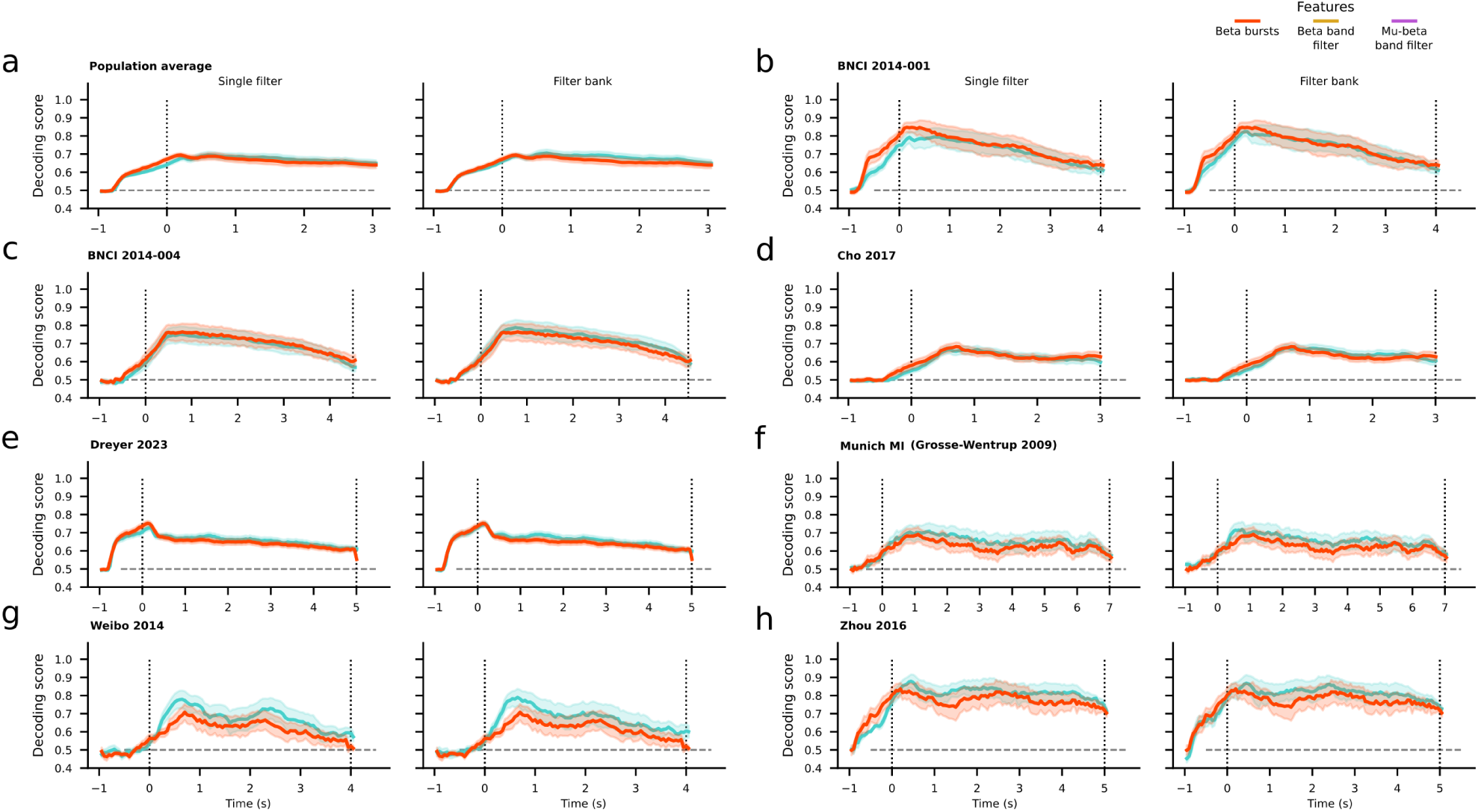
**(a)** Population average time-resolved decoding score and standard error for the beta burst convolution (red) and mu band (turquoise) filtering pipelines using a sliding window. Due to the different duration of the task for each dataset we restricted the time to the minimum trial period corresponding to 3 seconds. **(b – h)** Average, time-resolved decoding score and standard error per dataset of the same features using a sliding window. For each panel, the left subplot depicts the decoding results obtained using a single filter, while the right subplot depicts the results based on a filter bank technique. The beta burst results are the same for the pair of each panel. The horizontal dashed line corresponds to the expected chance level. Vertical dotted lines represent the onset and end of the trial period of each dataset. Horizontal lines on the top of each subplot show the results of pair-wise permutation cluster tests between the beta bursts and either filtering technique, with correction for multiple comparisons at significance level of 0.05. The color of the lines indicate which feature produces, on average, better results at any time point. A lack of color indicates no statistically significant differences between the compared features.

**Sup. Figure 4:**
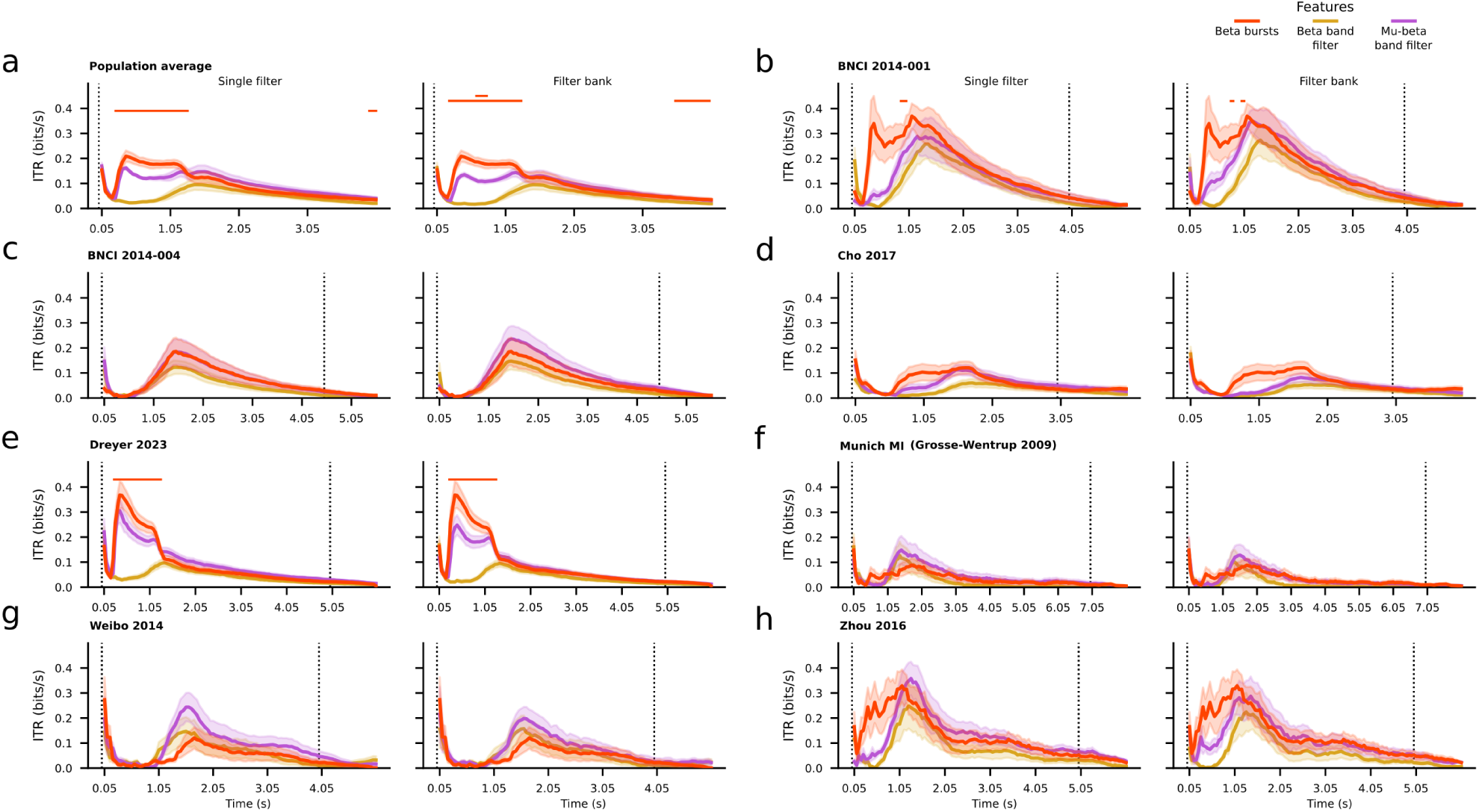
**(a)** Population average time-resolved information transfer rate (ITR) and standard error for the beta burst convolution (red), beta band (yellow) and mu-beta band (purple) filtering pipelines using a sliding window. Due to the different duration of the task for each dataset we restricted the time to the minimum trial period corresponding to 3 seconds. **(b – h)** Average, time-resolved information transfer rate (ITR) and standard error per dataset of the same features using a sliding window. For each panel, the left subplot depicts the ITR results obtained using a single filter, while the right subplot depicts the results based on a filter bank technique. The beta burst results are the same for the pair of each panel. Vertical dotted lines represent the onset and end of the trial period of each dataset. Horizontal lines on the top of each subplot show the results of pair-wise permutation cluster tests between the beta bursts and either filtering technique, with correction for multiple comparisons at significance level of 0.05. The color of the lines indicate which feature produces, on average, better results at any time point. A lack of color indicates no statistically significant differences between the compared features.

**Figure 5:**
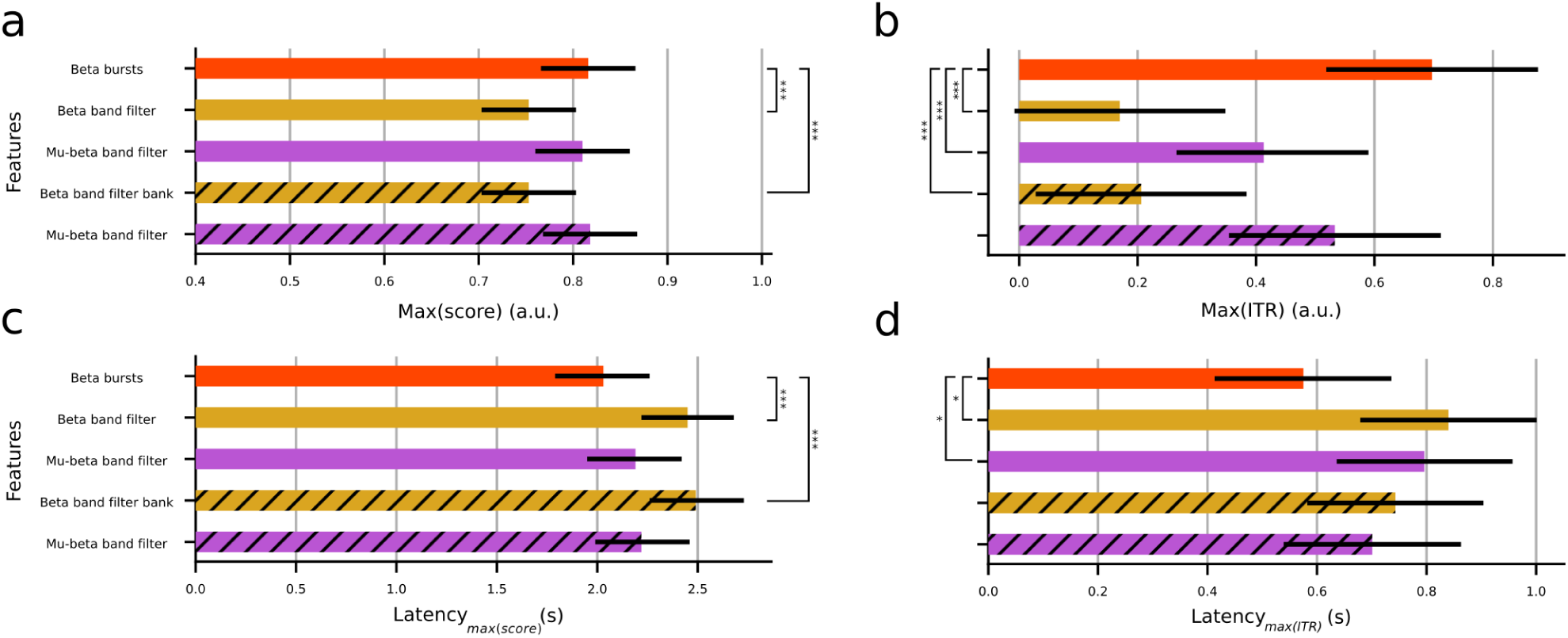
Population-level statistical analysis based on linear mixed models when using a sliding window per feature. **(a)** Average maximum decoding score. **(b)** Average maximum ITR. **(c)** Average latency to reach the maximum decoding score. **(d)** Average latency to reach the maximum ITR. Error bars show 95% confidence intervals. Hatches indicate the use of a filter bank technique. Asterisks indicate statistically significant differences among pairwise comparisons of the beta bursts and the rest of the features (* : p < 0.05. ** : p < 0.01, *** : p < 0.001). A lack of asterisks implies no statistically significant differences. Note that the log transform of the the maximum ITR and latency to maximum ITR were used for the statistical analysis (see Materials and Methods), but panels **b** and **d** depict results before applying the transformation for ease of comparisons with panels **a** and **c** respectively.

## Notes

### Competing Interest Statement

The authors have declared no competing interest.

